# Natural selection exerted by historical coronavirus epidemic(s): comparative genetic analysis in China Kadoorie Biobank and UK Biobank

**DOI:** 10.1101/2024.02.06.579075

**Authors:** Sam. C. Morris, Kuang Lin, Iona Y. Millwood, Canqing Yu, Jun Lv, Pei Pei, Liming Li, Dianjianyi Sun, George Davey Smith, Zhengming Chen, Robin Walters

## Abstract

**Background:** Pathogens have been one of the primary sources of natural selection affecting modern humans. The footprints of historical selection events – “selective sweeps” – can be detected in the genomes of present-day individuals. Previous analyses of 629 samples from the 1000 Genomes Project suggested that an ancient coronavirus epidemic ∼20,000 years ago drove multiple selective sweeps in the ancestors of present-day East Asians, but not in other worldwide populations.

**Results:** Using a much larger genetic dataset of 76,719 unrelated individuals from each of the China Kadoorie Biobank (CKB) and UK Biobank (UKB) to identify regions of long-range linkage disequilibrium, we further investigated signatures of past selective sweeps and how they reflect previous viral epidemics. Using independently-curated lists of human host proteins which interact physically or functionally with viruses (virus-interacting proteins; VIPs), we found enrichment in CKB for regions of long-range linkage disequilibrium at genes encoding VIPs for coronaviruses, but not DNA viruses. By contrast, we found no clear evidence for any VIP enrichment in UKB. These findings were supported by additional analyses using saltiLASSi, a selection-scan method robust to false positives caused by demographic events. By contrast, for GWAS signals for SARS-Cov2 susceptibility (critical illness, hospitalisation, and reported infection), there was no difference between UKB and CKB in the number located at or near signals of selection, as expected for a novel virus which has had no opportunity to impact the CKB/UKB study populations.

**Conclusions:** Together, these results provide evidence of selection events consistent with historical coronavirus epidemic(s) originating in East Asia. These results show how biobank-scale datasets and evolutionary genomics theory can provide insight into the study of past epidemics. The results also highlights how historic infectious diseases epidemics can shape the genetic architecture of present-day human populations.

## Background

Pathogens and their associated diseases have been widespread across human history (1). In particular, the transition from sparsely populated groups of hunter-gatherers to densely-packed farming communities in close vicinity to domesticated animals likely facilitated the spread of many novel pathogens from animals to humans, and then within and between human populations (2, 3). Despite widespread and substantial improvements in sanitation and treatment of infectious diseases, pathogens were still responsible for about a quarter of global deaths in 2019 (4). Thus, they are expected to have exerted substantial selective pressure on human populations throughout history; indeed, analysis of genetic data has suggested that pathogens represent the strongest selective effect on modern humans (5).

The impact of such past natural selection on the ancestors of modern humans can be observed in the genomes of present-day populations using a variety of statistical methods (e.g. Extended Haplotype Homozygosity (6), Population Branch Statistic (7), reviewed in (8)). These techniques have identified many immune-related loci inferred to have been targets of natural selection (9-12), supporting the hypothesis that pathogens play an important role in shaping patterns of human genetic variation. One such footprint of selection is known as a ‘selective sweep’: as an allele under positive selection rapidly increases in frequency within a population across generations, neighbouring alleles which are in linkage disequilibrium (LD) with the selected allele also rise to high frequency, erasing genetic diversity around the locus under selection (13, 14). Such selective sweeps can be detected by scanning the genome to identify e.g. long-range homozygous haplotypes (6, 9) or significant distortions of the haplotype frequency spectrum (15).

One set of likely pathogen-related targets of selection are virus interacting proteins (VIPs), which are classes of human proteins known to physically interact with or provide functions essential for replication of particular viruses. Previous analyses using sequencing data from the 1000 genomes project (16) identified an enrichment of selective sweep signals at genes encoding coronavirus (a type of RNA virus) VIPs in East Asian (EAS) but not European-ancestry (EUR) populations.

Conversely, no evidence for enrichment at genes for DNA-virus VIPs was found (12). Together, these results imply one or more historic coronavirus epidemics, either localised to East Asia or with signature(s) not detectable in EUR (e.g. due to different demographic histories or higher levels of post-selection genetic drift).

In the past few decades, there have been multiple epi/pandemics related to novel coronaviruses (i.e. COVID-19, MERS and SARS), which likely arose from zoonotic transmission. However, there also exist several ‘seasonal’ coronaviruses which are endemic in human populations, such as HCoV-229E and HCoV-NL63 (17). It is possible that these current seasonal coronaviruses originated as epidemics similar to the more recent epi/pandemics. Accordingly, the signals of selection identified by Souilmi et al (12) may reflect selection events related to the ancestors of these endemic viruses, and that there were multiple different sweeps related to several different endemic viruses. The COVID-19 pandemic predominantly affected older individuals in terms of mortality, suggesting that its selective impact at the population level through reproductive fitness may be limited. However, recent studies have indicated that long COVID, potentially among younger as well as older individuals, can lead to pathologies in various physical systems, including cardiovascular, neurological, cognitive and immune (18). Consequently, there may be a selective effect of long COVID mediated through its effect on these systems.

The finding that historical coronavirus epidemic(s) may have occurred in the ancestors of present-day EAS populations has important consequences for future studies on the effect of population-wide prior pathogen exposure on the risk of infection from novel diseases. Whilst the methodology employed by Souilmi, et al. gave statistically robust conclusions, their study was conducted on relatively small sample sizes (∼500 individuals across 5 EAS populations). To better characterise these historical selective sweeps, larger scale studies are needed, in both EAS and other populations; increasing sample size is known to improve precision when detecting weaker/more ancient sweeps (19, 20).

We have sought to replicate and extend the reported findings using a much larger genetic dataset comprising sets of 76,719 unrelated individuals from each of the China Kadoorie Biobank (CKB) and UK Biobank (UKB). We identified regions of long-range linkage disequilibrium (LRLD) in each population, and found that VIPs for coronaviruses, but not DNA viruses, were enriched for overlap with LRLD in CKB. By contrast, we found no clear evidence for any VIP enrichment in UKB. These findings were supported by concordant results for VIP enrichment at genomic regions identified by a selection scan using a different approach, in which distortion of the haplotype frequency spectrum was used to detect signals of selection. Together, these results provide further strong supporting evidence that one or more historical coronavirus epidemics occurred specifically in East Asia.

## Results

### Virus-interacting protein classification

VIPs are proteins expressed in humans that have been shown to interact with viruses, either physically or by providing functions essential for viral propagation. Genes encoding these proteins may be subject to selection driven by viral epidemics. We used a set of proteins grouped into VIP categories, as previously defined by Souilmi et al (12), based on low-throughput molecular methods and high-throughput mass-spectrometry (**Figure 1f)**, **Supplementary Table S1)**. VIPs were classified based on i) whether they primarily interact with DNA or RNA viruses; ii) whether or not RNA virus VIPs interact with coronaviruses; and iii) whether or not coronavirus VIPs interact with SARS-CoV-2 viruses. In addition, we defined a separate subset of 42 SARS-VIPs previously identified as being potentially sites of selection in past coronavirus epidemics (12) and which would be expected to be similarly identified in our analyses and which, therefore, could be used as a positive control to test the effectiveness of our analytical approach.

**Figure 1.**
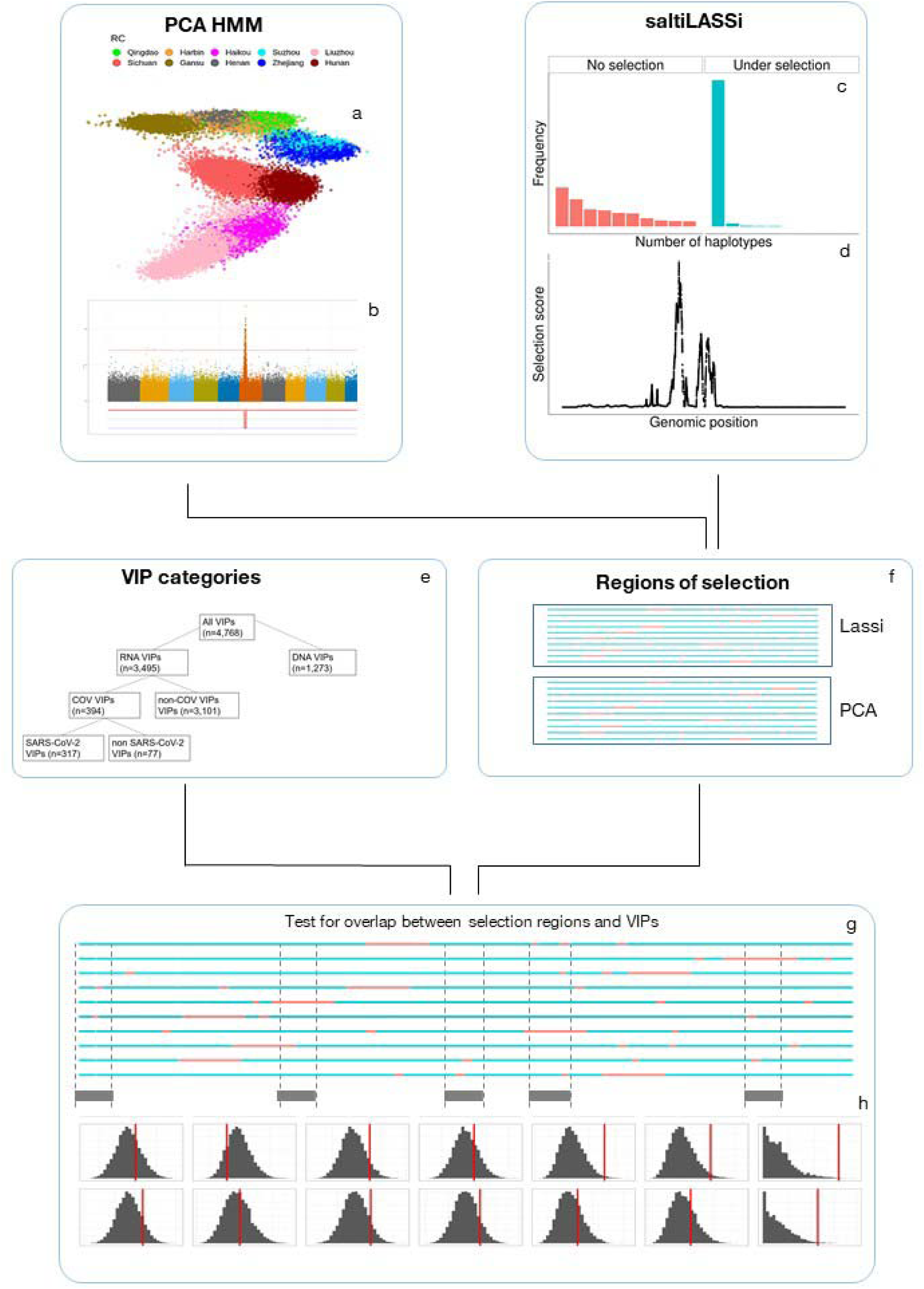
Schematic to estimate the enrichment of different VIP classes for regions of selection. We began by using two different methods to identify regions of as candidate regions of selection, PCA HMM and saltiLASSi. (a) The first step of the PCA HMM is to perform a PCA on the subset of unrelated individuals and then identify regions of the genome which show distortions in PC loadings using an HMM-based algorithm (b). Panel (b) shows a spike in the loadings of one principle component, caused by a region of long-range LD. The other method, saltiLASSi, identifies regions of the genome which show a strong distortion in the haplotype frequency spectrum in a sliding window approach (c) to calculate a selection scan test statistic. We then identified peaks of this test statistic (d). We then estimated the enrichment of each set of VIPS (e) with regions of selection (f). We calculated the empirical overlap between the regions of selection and different classes of VIPs (g) and then calculated whether this overlap is greater than expected by chance by permuting/bootstrapping the regions of selection across the genome to generate a null distribution (h).

### PCA-based identification of long-range LD regions

Natural selection and other demographic processes can result in regions of LRLD in the genome. In previous work, to facilitate genotype PCA analysis of the CKB cohort, we identified such regions of LRLD using an approach similar to one previously used in UKB (21), by applying an iterative hidden Markov-model based algorithm to principal components (PCs) derived from genotypes of 76,719 unrelated CKB individuals (see **Methods**) (22). Excluding the extended region of LD at the chromosome 6 HLA region (chr6:20-40Mbp), we identified 229 unique regions of LRLD (median length = 593.1Kbp, total length = 218.1Mbp) on the basis of distortions in the variant loadings of the top 11 PCs (i.e. those previously identified as being informative for geographic population structure in CKB) (**Supplementary Table S2**). Using the same approach for analysis of genotypes from a similar number of 76,719 randomly-selected unrelated white British individuals from UKB, applied to the top 5 (geographically-informative) PCs (23), we identified 326 LRLD regions (median length = 1070.0Kbp, total length = 518.77 Mbp) (**Supplementary Table S3**). Further sets of LRLD regions were defined based on splitting the CKB LDLR regions according to whether they were uniquely identified in CKB (n=128) or they overlapped with UKB LRLD regions (n=104) (**Supplementary Table S4-5**).

### Enrichment of long-range LD at VIP genes

We hypothesised that if a particular class of VIPs were the target of natural selection, then the genes encoding those VIPs would overlap with regions of LRLD more often than expected by chance. To test this, we compared the observed overlap of VIPs with LRLD regions with empirical null distributions, derived using sets of “decoy” LRLD regions generated by repeatedly redistributing the LRLD regions randomly across each chromosome while retaining their size characteristics, as illustrated in **Figure 1**. **Table 1** shows the results of this analysis for different classes of VIP and different sets of LRLD regions. Consistent results were found for 3 different methods for scoring LRLD - VIP overlap – i) any overlap, ii) >50% overlap, iii) total base-pair overlap (**Supplementary Tables S6-7**).

**Table 1.** Enrichment of regions of Long-range Linkage Disequilibrium (LRLD) at Virus Interacting Protein (VIP) genes. Enrichment and P-values (one-tailed test) were determined for the frequency with which the genes encoding different classes of VIPs overlap with sets of LDLR regions, compared with random permutation of LDLR locations across the genome. Number in brackets denotes the number of VIPs in that class included in the analysis. Enrichment and 95% CIs were derived from the median and 2.5% / 97.5% centiles of the null and P from the empirical 1-tailed test against the null. The HLA region (chr6:20-40Mb) and VIP genes lying within it were excluded from analysis. Asterisks next to P-values denote significance after multiple testing adjustment (P<0.05/(2*2.56), see Methods).

| LRLD Selection Region Set |  |  |  |  |  |  |  |  |
| --- | --- | --- | --- | --- | --- | --- | --- | --- |
| VIPs | CKB |  | UKB |  | CKB-only |  | CKB+UKB |  |
|  | enrichment | P | enrichment | P | enrichment | P | enrichment | P |
| All (4768) | 1.13 (0.98-1.31) | 0.047 | 0.98 (0.88-1.08) | 0.673 | 1.15 (0.95-1.42) | 0.076 | 1.11 (0.89-1.42) | 0.178 |
| RNA viruses (3495) | 1.19 (1.01-1.41) | 0.018 | 1.01 (0.90-1.14) | 0.429 | 1.25 (1.02-1.59) | 0.016 | 1.12 (0.87-1.48) | 0.183 |
| DNA viruses (1273) | 1.03 (0.83-1.28) | 0.419 | 0.92 (0.80-1.08) | 0.852 | 0.93 (0.70-1.31) | 0.687 | 1.12 (0.82-1.62) | 0.242 |
| CoV (394) | 1.50 (1.10-2.16) | 0.004* | 1.04 (0.84-1.32) | 0.374 | 1.89 (1.21-3.40) | 0.001* | 1.11 (0.74-2.00) | 0.316 |
| non-CoV (3101) | 1.16 (0.99-1.38) | 0.038 | 1.03 (0.91-1.16) | 0.337 | 1.20 (0.97-1.54) | 0.046 | 1.12 (0.86-1.48) | 0.195 |
| non-SARS (77) | 1.86 (1.00-4.33) | 0.021 | 1.07 (0.71-1.88) | 0.372 | 2.00 (1.00-8.00) | 0.023 | 1.67 (0.62-Inf) | 0.208 |
| SARS (317) | 1.41 (1.02-2.16) | 0.02 | 1.04 (0.82-1.34) | 0.41 | 1.86 (1.13-3.71) | 0.005* | 1.07 (0.65-2.14) | 0.424 |
| under-selection (42) | 2.50 (1.25-10.00) | 0.005* | 1.50 (0.86-4.00) | 0.083 | 2.00 (0.80-Inf) | 0.109 | 3.00 (1.20-Inf) | 0.011 |

Compared to the null distribution, there was strong evidence in CKB for LRLD enrichment at loci encoding the subset of SARS-VIPs (n=40 after exclusion of the HLA region) previously identified as likely sites of selection (enrichment ratio ER=2.50; 95% CI 1.25-10.00; P=0.005). This finding provides further population genetic evidence in support of the previous finding that one or more ancient coronavirus epidemics occurred in East Asia approximately 25,000 years ago, and indicates that, as expected, the identified regions of LRLD are enriched for signals of selection.

Using the same approach to test the other classes of VIPs, there was a strong signal in CKB that CoV-VIP genes (n=394) are enriched for regions of LRLD (enrichment ratio, ER=1.50; 95% CI 1.10-2.16; P=0.004) relative to the null. This LRLD enrichment was further investigated by classifying VIPs according to whether they have been found to be related to SARS viruses, or only related to other types of coronavirus (i.e. endemic coronaviruses). Both classes of VIPs displayed LRLD enrichment in CKB, with somewhat greater enrichment in non-SARS CoV-VIPs (ER=1.86, 95% CI 1.00-4.33; P=0.021) compared to SARS CoV-VIPs (ER=1.41, 95% CI 1.02-2.16; P=0.020). There was also suggestive evidence for more moderate enrichment of LRLD at genes encoding non-CoV-VIPs (ER=1.16; 95% CI 0.99-1.38; P=0.038), and for RNA-VIPs overall (ER=1.19; 95% CI 1.01-1.41; P=0.018). Conversely, DNA-VIPs (n=1,273) showed no enrichment for regions of LRLD (ER=1.03; 95% CI 0.83-1.28; P=0.419), again consistent with findings from the previous study by Souilmi et al (12).

By contrast with CKB, in UKB we found no evidence for enrichment of LRLD near to genes encoding CoV-VIPs (P=0.316), or for any other kind of VIPs (all P>0.05) (**Table 1)**. Furthermore, the LRLD enrichment observed for CKB was predominantly due to LRLD regions found only in CKB. For almost all classes of RNA VIP, enrichment for overlap with CKB-only LRLD was greater than for the main analysis while, conversely, overlap with LRLD regions identified in both CKB and UKB displayed less enrichment and/or was not statistically significant. The one exception was a 3-fold enrichment for the “under selection” SARS-CoV-VIP genes, although this was based on only 6 overlapping genes.

### Detecting signals of selection using saltiLASSi

In addition to selective sweeps, regions of LRLD may also arise from demographic processes such as population bottlenecks (24, 25), potentially confounding the above analysis. To address this issue, we aimed to replicate our findings using putative signals of natural selection identified using an unrelated method, saltiLASSi (15), which detects regions of the genome that display substantial distortions of the haplotype-frequency spectrum (illustrated in **Figure 1c**). Importantly, saltiLASSi is robust to false positives driven by e.g. demographic events. Applying saltiLASSi to the same sets of unrelated individuals from CKB and UKB, across the autosomes (again excluding the extended HLA region on chromosome 6), we identified 117 non-overlapping regions in CKB showing strong evidence of selection (median length = 175.2Kbp, total length = 35.3Mbp), and 118 regions in UKB (median length = 134.3Kbp, total length = 25.2Mbp) (**Supplementary Tables S8-9**. A total of 42 regions of selection overlapped between CKB and UKB, comprising 6.20Mbp of overlapping DNA.

### Enrichment of saltiLASSi regions at VIP genes

We assessed the proximity of VIP structural genes to these signals of selection, scoring for each VIP class the proportion lying within 10Kbp of a saltiLASSi-identified region (**Table 2**). For each of CKB and UKB, only 2.7% of DNA virus VIPs were close to at least one signal of selection, substantially fewer than for any other VIP class, consistent with the findings from our LRLD analysis and the previous finding of no evidence of selective sweep enrichment near DNA VIP genes (12). By comparison, in CKB the proportion of CoV-VIP genes in close proximity to a saltiLASSi signal of selection was substantially larger (27/394, 6.9%; P=5.6×10^-5^), with similar proportions for both SARS (6.9%; P=1.1×10^-4^) and non-SARS (6.5%; P=0.026) VIPs, in each case representing an enrichment of ∼2.5-fold relative to DNA virus VIPs. Further, for the SARs-VIPs previously identified as under selection, the proportion of genes close to a signal of selection was even larger (7/40, 17.5%; P=5.5×10^-8^), a 6.6-fold enrichment. In UKB, on the other hand, any enrichment of genes in proximity to signals of selection relative to DNA virus VIPs was much more limited. Nevertheless, there was near 2-fold enrichment for CoV-VIPs (20/394, 5.1%; 1.9-fold; P=0.012) and for SARs-VIPs (17/317, 5.4%; 2.0-fold; P=0.0096).

**Table 2.** Overlaps of genes encoding different classes of Virus Interacting Protein (VIP) with saltiLASSi selected regions in UKB and CKB participants. Genes encoding different classes of VIPs lie were scored according to whether they lay within 10Kbp of saltiLASSi-identified regions of selection (“overlap”), Enrichment and P-values were calculated relative to the proportional overlap for DNA virus VIPs using a two-sample proportion test, with the DNA VIP overlap as the null success rate. The HLA region (chr6:20-40Mb) and VIP genes lying within it were excluded from analysis. Asterisks next to P-values denote significance after multiple testing adjustment (P<0.05/(2*2.36)).

| saltiLASSi Selection Region Set |  |  |  |  |  |  |
| --- | --- | --- | --- | --- | --- | --- |
| VIP | CKB |  |  | UKB |  |  |
|  | Overlap | Enrichment | P | Overlap | Enrichment | P |
| DNA_ref | 34/1273 (2.7%) | 1.00 | ref | 35/1273 (2.7%) | 1.00 | ref |
| RNA | 144/3495 (4.1%) | 1.54 | 0.010* | 131/3495 (3.7%) | 1.36 | 0.048 |
| non-COV | 117/3101 (3.8%) | 1.41 | 0.035 | 111/3101 (3.6%) | 1.30 | 0.083 |
| CoV | 27/394 (6.9%) | 2.57 | 0.000* | 20/394 (5.1%) | 1.85 | 0.012* |
| non-SARS | 5/77 (6.5%) | 2.43 | 0.026 | 3/77 (3.9%) | 1.42 | 0.277 |
| SARS | 22/317 (6.9%) | 2.60 | 0.000* | 17/317 (5.4%) | 1.95 | 0.010* |
| under-selection | 7/40 (17.5%) | 6.55 | 0.000* | 2/40 (5.0%) | 1.82 | 0.199 |

To provide a more rigorous test for these observed enrichments of signals of selection near to VIPs, we adopted a bootstrapping approach similar to that used for LDLR. For each set of VIP structural genes, we scored the number that were in close proximity (within 10Kbp) to one or more saltiLASSi-identified regions, and compared this with a null distribution derived by redistributing the saltiLASSi-selected regions 10,000 times across the genome. In order to retain large-scale patterns of GC- and gene-content, the units of permutation were in the region of 300Kbp, so that it was necessary to exclude from analysis large saltiLASSi selection regions (those >500Kbp) and the VIPs in close proximity to them. Nevertheless, despite the resulting reduction in statistical power, in CKB both CoV-VIPs (ER = 2.12; 95% CI 1.13 – 5.67; P= 0.004) and SARS-VIPs (ER = 2.17; 95% CI 1.08 - 6.50; P=0.009) were once again strongly enriched for proximity to saltiLASSi-selected regions compared to the empirical null (**Table 3**). By contrast, we found no evidence for appreciable enrichment in any class of VIPs in UKB, consistent with the similar analysis of regions LRLD. These findings were robust to variations in the distance between the VIP structural gene and selection region used to define ‘proximity’, and to different sensitivity thresholds for detection of selection regions, with different sets of parameters giving qualitatively the same results (**Supplementary Tables S10a-d).**

**Table 3.**
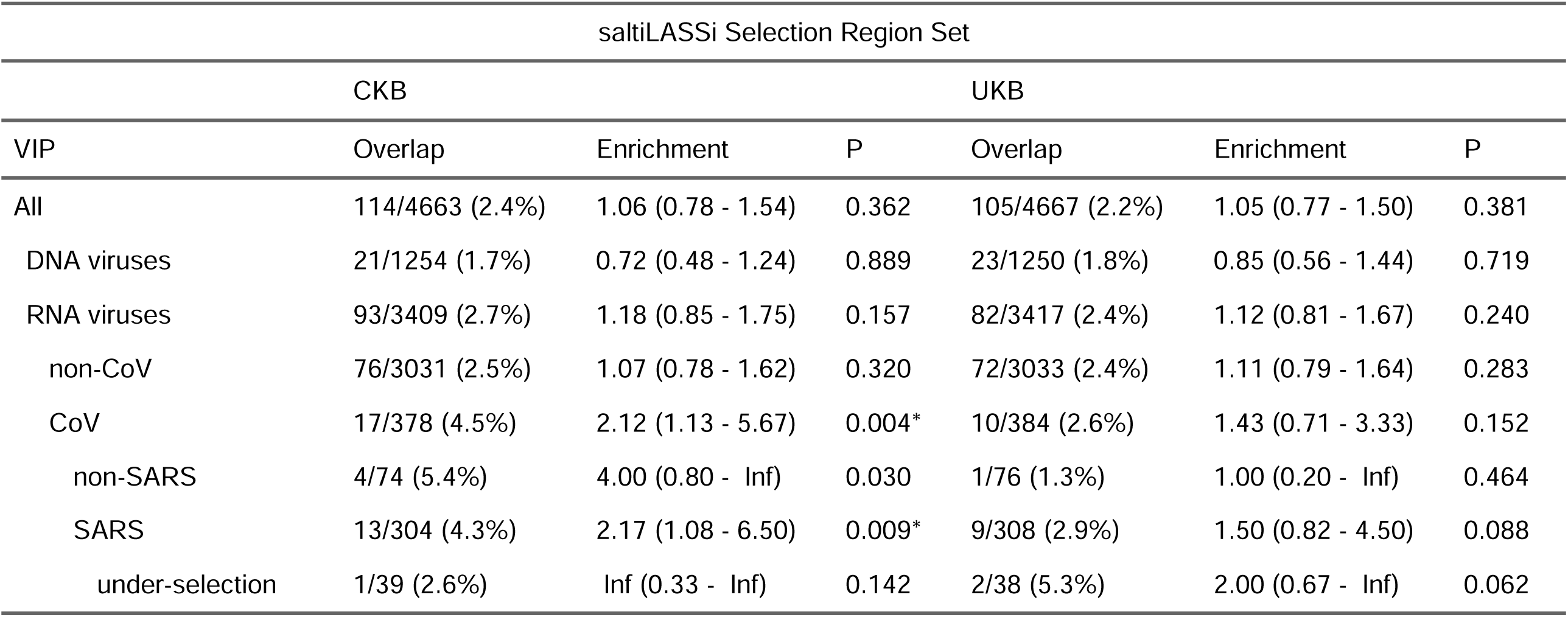
Overlaps between genes encoding different classes of Virus Interacting Protein (VIPs) and saltiLASSi selected regions in UKB and CKB participants. Enrichment and P-values (one-tailed test) were determined for the frequency with which the genes encoding different classes of VIPs lie within 10Kbp of saltiLASSi-identified regions of selection, compared with 10,000 bootstrap iterations of randomly distributing regions of selection across the genome, controlling for local gene density. Enrichment and 95% CIs are derived from the median and 2.5% / 97.5% centiles of the null distribution. The HLA region (chr6:20-40Mb) and other excludable regions (see methods), and VIP genes lying within them, were excluded from analysis. Asterisks next to P-values denote significance after multiple testing adjustment (P<0.05/(2*2.36).

### Regional analysis

Since CKB participants were recruited in 10 geographically diverse regions across China (26), we conducted further analyses to explore whether there were differences in selective signals between regions which might narrow down the geographical origins of the putative historical epidemic(s) which gave rise to the LRLD and saltiLASSi signals. Using phased CKB genetic data, we identified haplotypes which spanned the regions of LRLD that overlapped with COV VIPs (n=36) and determined the frequencies of a random subset of 2000 of the most common haplotypes in the different CKB recruitment regions (**Supplementary Table S11)**. No consistent pattern was discernible from this analysis, with no region showing strong evidence of having higher frequencies of these long-range haplotypes (**Figure 2a**). Similarly, we repeated saltiLASSi analyses for equal numbers of individuals from each CKB recruitment region (n = 10) and, for the 28 saltiLASSi selection signals which overlapped with COV-VIPs in the main analysis, scored the frequency with which these selection signals were identified when restricting analysis to individuals from a single recruitment centre (**Figure 2b**. Again, no clear difference was observed between regions, with 78-92% (mean 87%) of the signals being replicated across each region.

**Figure 2.**
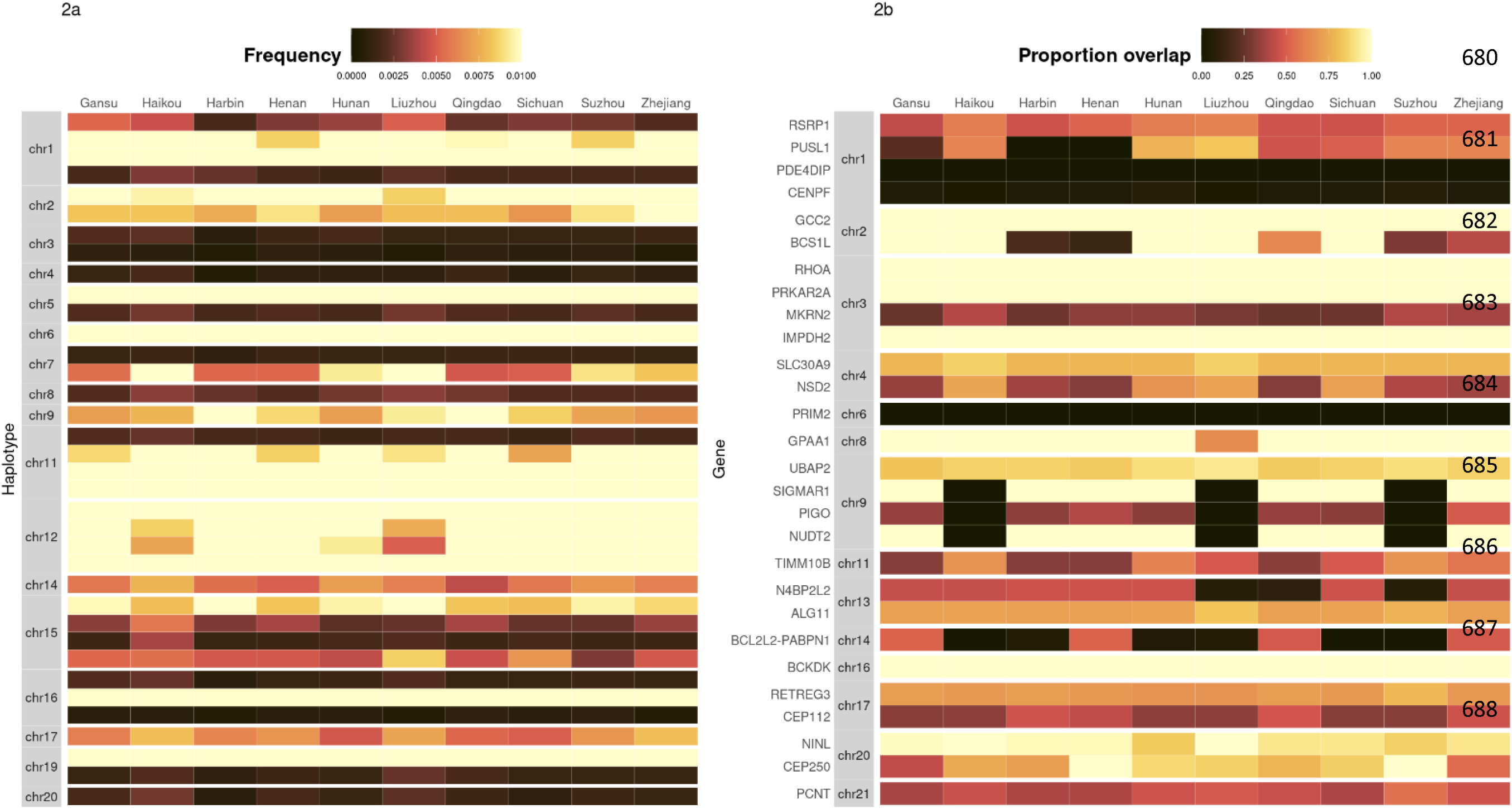
b) **Frequency, in each recruitment region (n=10), of the most common haplotypes which span the regions of long-range LD which overlap with the COV-VIPs (n=36).** We took the phased CKB data and extracted the haplotypes which covered the regions inferred to be under selection by the LRLD selection method and calculated the frequency of the most common haplotype (across the entire dataset), within each region. Each region was subsamples to 2000 randomly selected individuals to ensure comparability across regions. b) **Replication of saltiLASSi selection hits in different recruitment centres (RC) in China (n=10).** We analysed each recruitment region using saltiLASSi separately, after subsampling the number of individuals to match the RC with the fewest number of individuals and determined whether selection hits obtained from analysing the full cohort were replicated in each RC. The colour of each tile represents the proportion of the selection hit inferred in the full cohort which was covered by the selection hit inferred a given CKB region.

### Overlap with SARS-Cov2 susceptibility GWAS associations

To explore whether signals of selection overlapped with recently published GWAS associations for COVID susceptibility (27), we selected all 51 lead SNPs which reached genome-wide significance (i.e. 5 x 10^-8^) for “Critical illness", "Hospitalized" and "Reported infection" and counted the overlap between these loci (defined by the region within 200Kbp of the lead variant) and either the LRLD or the saltiLASSi regions identified in CKB and UKB, again using permutation across the genome to provide an empirical null. Of the 51 lead variants for COVID-19 susceptibility, 6 overlapped with regions of LRLD in CKB (ER=1.67, 95% CI 0.71 – Inf, P=0.257), and 5 overlapped with UKB LRLD regions (ER=0.71, 95% CI 2-12, P=0.836). The corresponding analysis using saltiLASSi regions, and the bootstrapped regions from the previous section, yielded similar results for both CKB (7/51; ER=1.75, 95% CI 0.78 – 7.00, P=0.09) and UKB (5/51; ER=1.25, 95% CI 0.56 – 5.00 P=0.27). The saltiLASSi results were robust to variation in the size of the window surrounding the GWAS lead variants (50Kbp-500Kbp) (**Supplementary Table S12**).

## Discussion

Viral epidemics are expected to exert relatively fast-acting selection on the human genome. Such events can leave footprints in the form of ‘selective sweeps’, in which linked neutral variants ‘hitchhike’ to higher frequency, thereby reducing genetic variation around a selected locus and generating regions of LRLD (13, 28). Previous work provided evidence that an ancient coronavirus epidemic(s) more than 20,000 years ago drove selective sweeps(s) in the genomes of EAS individuals (12). However, this study was based on a relatively small size of ∼100 individuals from each of 5 populations, and simulations have shown progressive increases in sweep detection accuracy with increasing sample sizes; for instance, for a sweep occurring 1000 generations ago, there is a 1.6 fold increase in power when increasing the number of haplotypes from 10 to 50 (20). We sought to further investigate these potential signatures of historic viral epidemics in a substantially larger dataset comprising ∼70k individuals from each of CKB and UKB, and again found that genes encoding proteins which interact with coronaviruses are significantly more likely to be near regions of selection, in Chinese but not British individuals.

We first assessed enrichment of VIP genes in regions of LRLD, relative to a null distribution in which regions of the same size were repeatedly randomly distributed around the genome, an approach widely used to assess the significance of associations of one genomic feature with another (e.g. (29, 30)). DNA virus VIP genes showed no enrichment of LRLD, consistent with the previous finding of no signals of selection at these VIPs. By contrast, genes encoding a set of SARS-VIPs that were previously identified as potentially under selection showed a 2.5-fold enrichment of nearby LRLD regions. Together, these findings indicate that this approach is both well-calibrated and capable of detecting signatures of selection.

However, LRLD does not arise exclusively from selective sweeps and can arise due to various non-selective processes, such as restrictions on genetic recombination due to genomic structural variants such as inversions (31-33). Furthermore, demographic events such as population bottlenecks can produce regions of LDLR (24) and other signatures which mimic positive selection and can thus confound selection scans. Such processes could influence both the previous analyses (12) and our analysis based on LRLD. To address these potential issues, therefore, we conducted a separate analysis in which signals of selection were instead identified using saltiLASSi, which accounts for the spatial distribution of the sweep test statistic across the genome and is thereby more robust to non-selection demographic events. Once again, there was no enrichment of signals of selection near to DNA virus VIPs, while the previously identified set of SARS-VIPs potentially under selection showed substantial enrichment relative to other VIP classes.

These two analyses, based on methodologically distinct approaches to identification of regions of selection, gave consistent results when applied to different classes of VIP. Both identified strong enrichment of selection regions in CKB near to CoV-VIPs and SARS-VIPs, but not near non-CoV-VIPs. Both methods also identified enrichment at non-SARS-VIPs (comparable to that at SARS-VIPs), although this was based on a smaller number of VIPS in this class, giving wide confidence intervals. Conversely, we found no evidence of any significant enrichment of any classes of VIPs in UKB. Furthermore, the observed enrichment in CKB was almost entirely derived from regions of selection identified only in CKB and not in UKB. These findings, that there is enrichment of signatures of selection at genes encoding CoV-VIPs in CKB but not UKB, are entirely consistent with the hypothesis that one or more historical epidemics of coronaviruses (or other viruses which interact similarly with cellular processes) occurred in the ancestors of modern-day EAS populations.

Given the geographical restriction of the putative selective sweep(s), it was of interest to explore whether the much larger sample size in our analyses enabled any greater geographical resolution of the origins of such sweep(s). However, we found no clear evidence that enabled localisation of the enrichment of selection signals to one or more particular regions in China. This is perhaps not surprising, as many population migrations and population mixing have taken place across China in the past 20,000 years which are likely to have obscured any region-specific signals (34). Alternatively, any epidemic may have been widespread across East Asia, which would be consistent with the results of Souilmi et al who found signals in other East/South East Asian countries.

It is known that environmental pressures in the history of a population may confer lasting adaptations to humans (35-37). Therefore, it is plausible that widespread, and potentially repeated, historical coronavirus epidemics in East Asia may have provided a degree of resistance for modern-day East Asians to the recent COVID-19 pandemic. Evidence has shown that different populations have different mortality risks from severe COVID-19 (38, 39). Whilst it is clear that sociodemographic factors and provision of appropriate health care play a substantial part in these differences, there is also the possibility that variants with protective effects against COVID-19 may be distributed differentially across populations. For instance, African American ancestry has been reported as an independent risk factor for hospitalisation from COVID-19 (40), If ancient and current coronavirus epidemics have VIPS in common, the putative historic epidemic in East Asia may have driven selective sweeps in regions of the genome which are currently under selection by COVID-19. This could be manifested in a higher number of overlaps between GWAS hits and regions of selection in Chinese compared to British individuals. However, we found no discernible differences between the populations. Whilst this may be due to a low number of GWAS hits, or may reflect insufficient EAS individuals in GWAS of COVID-19 susceptibility, this may also point to differences in the proteins relevant to present-day and ancient coronaviruses. A further consideration is that signatures of selection from the putative historical epidemic(s) will included contributions from the long-term effects of the virus, analogous to the long-term “long Covid” effects of SARS-Cov2, which are not included in GWAS of COVID-19 susceptibility.

Nevertheless, observational evidence suggests that countries in East Asia have a substantially lower acute case-fatality rate than comparable countries in Western Europe (41). While it is very likely a substantial part of the observed differences in severity and case-fatality rates between different populations are due to non-biological factors, e.g. public policy and differences in social behaviour, these data, alongside VIP enrichment results from this and previous studies, suggest that EAS populations may have a higher frequency of alleles protective against severe COVID-19, that may be one cause of the reduced case-fatality rates in these populations. For instance, non-synonymous mutations in *TMPRSS2* that confer decreased COVID-19 susceptibility are found at higher frequencies in EAS (36%) than EUR (23%) (42).

Variation in genetic susceptibility to disease across different ancestries, driven by differing nature selection environments, is well documented; for instance, alleles which provide a protective effect against Malaria are found at substantially higher frequencies in West Africa than in Europe (43, 44). Further, *in vivo* studies have shown that the transcriptional response of primary macrophages to live bacterial pathogens varies between ethnicities, and that genetic effects in the immune response are strongly enriched for recent, population-specific signatures of adaptation (45). It is also known that epi/pandemics drive population-level immunity which protects against future outbreaks; historical evidence cites that during the initial outbreak of the plague across Europe, during 1347 to 1353, not a single town was re-infected two or more years running (46). Thus, it is plausible that past coronavirus selective sweeps in EAS populations have provided a degree of resistance to COVID-19. For instance, the *ACE2* locus on the X chromosome (not included in our analysis, which was restricted to autosomes) has reduced haplotype diversity in EAS, one of the two major haplotypes being associated with appreciably lower SAR-Cov2 severity (47). Furthermore, single cell transcription analyses have shown that natural selection has driven population-specific differences in the immune response to SARS-Cov2 (48).

The key strength of our study is the use of biobank-scale data to provide greater power to identify signals of selection. Apart from the large sample sizes, the genetic data in CKB and UKB were generated using similar Axiom arrays with 50% of the genotyped variants being the same, reducing the chance of possible array-specific confounding. Moreover, we used a method, saltiLASSi, which is more robust to false-positives than the approach used by Souilmi et al (15), and the involvement of 10 geographically diverse regions in CKB enabled us to explore possible regional variations within China. However, the study also has limitations, most notably that, unlike the study by Souilmi et al. which was based on sequence data, we used genotype array data. This means we could not explore more detailed parameters of putative sweeps, such as strength and age.

## Conclusions

This study provides further evidence that historical coronavirus epidemic have shaped the genetic landscape of East Asian populations, as observed through significant enrichment of coronavirus interacting protein (CoV-VIP) genes in regions undergoing selection. Our findings, leveraging large biobank-scale datasets, reinforce the important role of pathogen epidemics in human evolutionary history but also underscore the potential influence of ancestral viral exposures on population-specific disease susceptibility as a research avenue.

## Declarations

### Ethics approval and consent to participate

Approval of the study was obtained from ethics committees or institutional review boards at the University of Oxford, the Chinese Center for Disease Control and Prevention (China CDC), the Chinese Academy of Medical Sciences, and all participating regions.

### Consent for publication

This research was funded in whole, or in part, by the Wellcome Trust [212946/Z/18/Z, 202922/Z/16/Z, 104085/Z/14/Z, 088158/Z/09/Z]. For the purpose of Open Access, the author has applied a CC-BY public copyright licence to any Author Accepted Manuscript version arising from this submission

### Availability of data and materials

The datasets supporting the conclusions of this article are included within the article and its additional files. Sharing of genotyping data is currently constrained by the Administrative Regulations on Human Genetic Resources of the People’s Republic of China. Access to these and certain other data is available through collaboration with CKB researchers.

### Authors’ contributions

Conceived and designed the study: **RGW, SCM, KL, GDS, ZC**; performed analyses: **SCM, KL**; interpreted results: **RGW**, **KL**, **SCM**, **ZC**, **GDS**; and wrote the manuscript: **RGW**, **SCM**, **ZC**. Provided administrative, technical or research support **CY**, **IYM**, **JL**, **PP**, **LL**, **DS**

## Acknowledgements

The most important acknowledgement is to the participants in the study and the members of the survey teams in each of the 10 regional centres. We also thank the project development and management teams based at Beijing, Oxford, and the 10 regional centres.

## Methods

### Statistics and reproducibility

The details of each analysis are outlined in the methods section and all of the code has been made publicly available on GitHub at https://github.com/sahwa/CKB_COVID_selection

### Study populations and genotyping data

#### CKB

China Kadoorie Biobank is a population based prospective cohort of >512,000 participants, of whom 100,706 had available genotyping data as previously described (22, 26). Individuals were genotyped on custom-designed Axiom® arrays optimised for individuals with East Asian ancestry, on which 340,562 genotyped variants overlapped with the UK Biobank genotype array. Analyses were based on 513,164 variants passing quality control on both array versions and in all genotyping batches. One individual from each pair of individuals with KING kingship coefficient cutoff >0.05 (determined using an LD-pruned set of 171,236 variants) was removed to create a set of 76,719 unrelated individuals used in the present study.

#### UKB

Genotyping data for 805,426 directly-genotyped variants in UKB participants was available under project 50474. We selected self-identified ‘White British’ individuals based on Data-Field 22006 and used an LD-pruned set of 230,948 variants to define an unrelated set of individuals using KING kingship coefficient cutoff >0.05. From the set of 348,845 unrelated individuals, we randomly selected 76,719 samples to match the number of CKB samples.

### Virus Interacting Proteins

VIPs (n=4,768 after exclusions) and their categorisations were as defined by Soulimi et al 2021 (12), with genomic coordinates of structural genes (build 37) as downloaded using Ensembl v102 (49). VIPs were excluded whose genes were non-autosomal or which lay within an extended MHC region (chr6:21,745,208-39,042,510) defined based on results from LDLR identification in CKB. Similarly, for all analyses we only considered VIPs which overlapped with regions genotyped in the CKB and UKB datasets, by splitting the genome up into regions of 500Kb non overlapping segments and then only considering VIPs which are fully covered by a segment.

### Identification of putative regions under selection

#### a) Long-range linkage disequilibrium

The method for identification of LRLD regions as applied to CKB has been described previously (49). Adapted from an approach to remove distortions principal components analysis (PCA) (23), we conducted a systematic iterative search for regions of LRLD by applying a hidden Markov model (HMM) to PCA loadings. For each biobank, an initial variant set was derived by filtering to remove variants with MAF<0.01 and Hardy-Weinberg P<10^-4^. We also performed local pairwise LD pruning using plink --indep-pairwise 50 5 0.2 (50). We then performed PCA of the pruned genotypes using flashpca (51). Starting with the variant loadings for PC1, and for each chromosome in turn, variants were assigned to one of two states: under selection (SR) or not, using a hidden Markov model. The emission probability of a variant being within a SR region, given its absolute loading value, was determined from the cumulative p-value from the chi-squared distribution with one degree of freedom. The transition probability between the states is in proportion to EAS recombination rates (downloaded from SniPA (52)); over a scaling factor of 1E+7. The loadings were decoded using the forward-backward algorithm given by Rabiner (53), and variants with a marginal likelihood >0.5 were assigned to the final set of selected regions. SNPs were assigned to one of the two states. Regions were defined by combining consecutive SNPs of the same states, while borders are at the middle points of two consecutive SNPs of different states.

In the next iteration, the SNPs covered by the SR regions were removed and PCA was performed again. Then the newly identified SR regions were merged with the previous sets. Once the detection of SR set converged, with no additional SR regions to be discovered, the number of PCs to be parsed were incremented by 1. In total we analysed the loadings of the first 11 and 5 PCs for CKB and UKB, respectively, these being the PCs informative for geographical population stratification.

In addition to the CKB and UKB SR sets, we also defined sets of selection regions which were i) the intersection of CKB and UKB or ii) found in CKB but not in UKB.

#### b) PCA loadings permutation test

To test whether the overlap between VIPs and SR regions was greater than would be expected by chance, we used bedtools (version v2.30.0) (54) to generate decoy SR sets, to enable derivation of empirical P values. Given a SR set, for each chromosome, the locations of the selection regions were randomly shuffled, with no overlaps, 10,000 times. We collected the corresponding 10,000 “decoy” selection region sets.

Adding 10Kb upstream and downstream to each VIP, the overlap between a VIP gene set and a SR set was compared with the overlap in the decoy SR sets, to give empirical p-values for three sets of features: the number of VIP genes overlapping selection regions by at least 1bp; the number of VIP genes with greater than half covered by selection regions; and the number of base-pairs covered by the regions. The rank of the genuine overlapping statistics, among the sorted 10,000 decoy values, was taken as the empirical P-value.

### VIP set Multiple Testing Correction

To account for testing multiple sets of sometimes correlated VIPs, we applied the procedure from Machado 2007 (55) to determine a Bonferonni correction to apply to the P-value threshold. To derive, *S*, the approximate number of independent tests, first let *M* be an *n* * *p* matrix, where *n* is the number of VIP sets and *P* the total number of VIPs across all sets. The elements of *M*, denoted by *M*_ij_, are defined as follows:

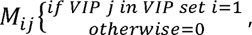 where *i* = 1,2 ….,*n* and *j* = 1,2 ….. *p*. Let *v*_1_, *v*_2_,…., *v_n_* be the eigenvalues of *M*. Rescale all eigenvalues so that they sum to n: 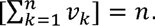 For each eigenvalue *v_k_*, modify such that *v_k_* = min (*v_k_*, 1). The sum of the modified eigenvalues, 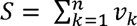 gives the number of approximately independent tests, and we accordingly use 0.05/S as our significance threshold.

### LASSi

Genotype data from each biobank were phased using shapeit v4.1.3 using default settings (56). The saltiLASSi (v-1.1.1) (15) algorithm was applied to the same CKB and UKB datasets of 76,719 participants each. After initial QC (22) no allele frequency/count filters were applied to the genotype data before applying the selection scan. We used the settings --winsize 10 and --winstep 1, with all other parameters as default. A small window size was selected to give increased power to detect relatively old or weak selective sweeps (15). The value *L*, the saltiLASSI composite-likelihood ratio test statistic, was used as a metric for the strength of evidence for a selective sweep and the basis on which to define a region under selection.

“Selected regions” (SRs) were defined as regions of contiguous SNPs which had *L* values above the 0.99 quantile for all *L* values for that chromosome and at least 200 SNPs away from another contiguous region of SNPs above the 0.99 quantile.

### Bootstrapping saltiLASSi regions of selection

To determine whether the overlap between the regions of selection identified by saltiLASSi and different classes of VIPs was greater than would be expected by chance, we used the bootRanges function from the nullRanges R library (57). Following the steps in the vignette, we used the EnsDb.Hsapiens.v86 genome (58) and excluded the following regions

i. hg38.Kundaje.GRCh38_unified_Excludable
ii. hg38.UCSC.centromere
iii. hg38.UCSC.telomere
iv. hg38.UCSC.short_arm
v. the extended HLA region (chr6:21,745,208-39,042,510)
vi. MT, chrY and chrX

The length of isochores (i.e. regions which capture large-scale patterns of GC and gene content) across the human genome are in the range of 300Kb - 1Mb (59). Hence, in order to capture the structure of the isochores, we also removed any regions of selection which were longer than 500Kb.

We segmented the remaining genome according to gene density. We performed 10,000 bootstrap iterations and calculated the overlap between each VIP set and the bootstrapped saltiLASSi selection regions. The empirical P-value was given by the proportion of times the randomly permuted selection regions had a greater number of overlaps with the VIP set than the true number of selection regions, divided by the number of bootstrap iterations. 97.5% enrichment intervals around the enrichment ratios were obtained by dividing the true proportion of overlaps by the 97.5 quantiles of the bootstrapped distribution of overlaps. We applied the same P-value threshold adjustment as in the LRLD analysis (**VIP set Multiple Testing Correction).**

### Relative enrichment of VIP sets relative to DNA VIPs

Our analysis and that of Souilmi et al (2019) suggested that DNA VIPs are not under any kind of detectable selection. Therefore, we used the overlap between saltiLASSi selection regions and DNA VIPs as a null success rate with which to compare other VIP sets against. We calculated P-values of the relative enrichment of non-DNA VIP sets relative to DNA VIP sets using stats::prop.test in R (v4.3.2).

### Overlap between selection regions and GWAS hits

We downloaded the lead hits for 3 different COVID-19 related traits (“Critical illness", "Hospitalized" and "Reported infection"), from https://app.covid19hg.org/, retaining only hits in autosomal regions. We also removed any hits which fell inside the extended HLA region (∼chr6:20-40Mb). Due to the small overall number of hits, to maximise power to detect any signals, we combined the lead hits for all 3 traits together, resulting in a total of 51 loci.

We counted the number of overlaps between each class of lead hit loci (defined by the region within 200Kbp of the lead variant) and either the i) regions of long-range linkage disequilibrium (LRLD) or ii) saltiLASSi regions of selection. We also tested varying the window added around each GWAS loci between 50Kb, 100Kb, 200Kb and 500Kb. To determine whether the overlap between GWAS loci and selection regions was greater than expected by chance, we used the same bootstrapping procedure as in the previous section, using bootranges for the saltiLASSi regions and the decoy regions for regions of LRLD.

