## Supplementary tables for "Natural selection exerted by historical coronavirus epidemic(s): comparative genetic analysis in China Kadoorie Biobank and UK Biobank": COVID_selection_suppinfo_tables.docx

Supplementary Table S10a. Counts of overlaps (defined as being within 5Kbp of the Virus Interacting Protein (VIP) structural gene) between different classes of VIPs and saltiLASSi selected regions in the UKB and CKB participants. Enrichment and P-values (one-tailed test) were determined for the frequency with which the genes encoding different classes of VIPs overlap with saltiLASSi-identified regions of selection, compared with 10,000 bootstrap iterations of randomly distributing regions of selection across the genome, controlling for local gene density. The HLA region (chr6:20-40Mb) and other excludable regions (see methods) and VIP genes lying within it were excluded from analysis. Asterisks next to P-values denote significance after multiple testing adjustment (P<0.05/(2*2.36).

| Selection Region Set (5Kbp) | | | | | | |
| --- | --- | --- | --- | --- | --- | --- |
|  | CKB |  |  | UKB |  |  |
| VIP | Overlap | Enrichment | P | Overlap | Enrichment | P |
| All | 111/4768 (2.3%) | 1.07 (0.78 - 1.56) | 0.341 | 102/4768 (2.1%) | 1.06 (0.78 - 1.52) | 0.344 |
| DNA viruses | 21/1273 (1.6%) | 0.75 (0.49 - 1.31) | 0.858 | 23/1273 (1.8%) | 0.88 (0.58 - 1.53) | 0.657 |
| RNA viruses | 90/3495 (2.6%) | 1.18 (0.86 - 1.80) | 0.157 | 79/3495 (2.3%) | 1.13 (0.81 - 1.68) | 0.225 |
| non-COV | 74/3101 (2.4%) | 1.09 (0.78 - 1.64) | 0.306 | 69/3101 (2.2%) | 1.10 (0.78 - 1.68) | 0.280 |
| COV | 16/394 (4.1%) | 2.00 (1.14 - 5.33) | 0.005* | 10/394 (2.5%) | 1.43 (0.77 - 5.00) | 0.126 |
| non-SARS | 3/77 (3.9%) | 3.00 (0.60 - Inf) | 0.085 | 1/77 (1.3%) | 1.00 (0.25 - Inf) | 0.440 |
| SARS | 13/317 (4.1%) | 2.17 (1.08 - 6.50) | 0.007* | 9/317 (2.8%) | 1.80 (0.82 - 9.00) | 0.072 |
| under-selection | 1/40 (2.5%) | Inf (0.33 - Inf) | 0.133 | 2/40 (5.0%) | 2.00 (0.67 - Inf) | 0.056 |

Supplementary Table S10b. Counts of overlaps (defined as being within 20Kbp of the Virus Interacting Protein (VIP) structural gene) between different classes of VIPs and saltiLASSi selected regions in the UKB and CKB participants. Enrichment and P-values (one-tailed test) were determined for the frequency with which the genes encoding different classes of VIPs overlap with saltiLASSi-identified regions of selection, compared with 10,000 bootstrap iterations of randomly distributing regions of selection across the genome, controlling for local gene density. The HLA region (chr6:20-40Mb) and other excludable regions (see methods) and VIP genes lying within it were excluded from analysis. Asterisks next to P-values denote significance after multiple testing adjustment (P<0.05/(2*2.36).

| Selection Region Set (20Kbp) | | | | | | |
| --- | --- | --- | --- | --- | --- | --- |
|  | CKB |  |  | UKB |  |  |
| VIP | Overlap | Enrichment | P | Overlap | Enrichment | P |
| All | 118/4768 (2.5%) | 1.03 (0.76 - 1.49) | 0.434 | 113/4768 (2.4%) | 1.04 (0.77 - 1.47) | 0.405 |
| DNA viruses | 21/1273 (1.6%) | 0.68 (0.45 - 1.17) | 0.929 | 26/1273 (2.0%) | 0.87 (0.59 - 1.44) | 0.679 |
| RNA viruses | 97/3495 (2.8%) | 1.15 (0.84 - 1.70) | 0.186 | 87/3495 (2.5%) | 1.10 (0.80 - 1.61) | 0.283 |
| non-COV | 80/3101 (2.6%) | 1.07 (0.77 - 1.60) | 0.352 | 77/3101 (2.5%) | 1.08 (0.78 - 1.60) | 0.313 |
| COV | 17/394 (4.3%) | 2.12 (1.06 - 5.67) | 0.007* | 10/394 (2.5%) | 1.25 (0.67 - 3.33) | 0.217 |
| non-SARS | 4/77 (5.2%) | 2.00 (0.80 - Inf) | 0.038 | 1/77 (1.3%) | 0.50 (0.20 - Inf) | 0.511 |
| SARS | 13/317 (4.1%) | 1.86 (1.00 - 6.50) | 0.016* | 9/317 (2.8%) | 1.50 (0.75 - 4.50) | 0.127 |
| under-selection | 1/40 (2.5%) | Inf (0.33 - Inf) | 0.162 | 2/40 (5.0%) | 2.00 (0.67 - Inf) | 0.074 |

Supplementary Table S10c. Counts of overlaps (defined as being within 50Kbp of the Virus Interacting Protein (VIP) structural gene) between different classes of VIPs and saltiLASSi selected regions in the UKB and CKB participants. Enrichment and P-values (one-tailed test) were determined for the frequency with which the genes encoding different classes of VIPs overlap with saltiLASSi-identified regions of selection, compared with 10,000 bootstrap iterations of randomly distributing regions of selection across the genome, controlling for local gene density. The HLA region (chr6:20-40Mb) and other excludable regions (see methods) and VIP genes lying within it were excluded from analysis. Asterisks next to P-values denote significance after multiple testing adjustment (P<0.05/(2*2.36).

| Selection Region Set (50Kbp) | | | | | | |
| --- | --- | --- | --- | --- | --- | --- |
|  | CKB |  |  | UKB |  |  |
| VIP | Overlap | Enrichment | P | Overlap | Enrichment | P |
| All | 140/4768 (2.9%) | 1.01 (0.76 - 1.44) | 0.451 | 140/4768 (2.9%) | 1.03 (0.77 - 1.43) | 0.412 |
| DNA viruses | 27/1273 (2.1%) | 0.73 (0.49 - 1.23) | 0.901 | 33/1273 (2.6%) | 0.89 (0.61 - 1.43) | 0.667 |
| RNA viruses | 113/3495 (3.2%) | 1.12 (0.83 - 1.64) | 0.220 | 107/3495 (3.1%) | 1.08 (0.80 - 1.55) | 0.302 |
| non-COV | 95/3101 (3.1%) | 1.06 (0.77 - 1.53) | 0.363 | 95/3101 (3.1%) | 1.07 (0.79 - 1.53) | 0.321 |
| COV | 18/394 (4.6%) | 1.80 (1.00 - 4.50) | 0.017* | 12/394 (3.0%) | 1.20 (0.71 - 3.00) | 0.239 |
| non-SARS | 4/77 (5.2%) | 2.00 (0.67 - Inf) | 0.074 | 2/77 (2.6%) | 1.00 (0.33 - Inf) | 0.363 |
| SARS | 14/317 (4.4%) | 1.75 (0.93 - 4.67) | 0.026 | 10/317 (3.2%) | 1.25 (0.71 - 3.33) | 0.195 |
| under-selection | 1/40 (2.5%) | 1.00 (0.33 - Inf) | 0.212 | 3/40 (7.5%) | 3.00 (0.75 - Inf) | 0.035 |

Supplementary Table S12. Overlaps between regions of selection and 51 GWAS hits. Enrichment and P-values (permutation test) were determined for the frequency with which the GWAS hits for COVID lie within different distances (Window) of regions of LASSi-identified regions of selection, compared with 10,000 bootstrap iterations of randomly distributing regions of selection across the genome, controlling for local gene density. Enrichment and 95% CIs are derived from the median and 2.5% / 97.5% centiles of the null distribution.

| Population | Window | Overlap | Median enrichment | P |
| --- | --- | --- | --- | --- |
| CKB | 50Kb | 6 (11.8%) | 2.00 (0.86- Inf) | 0.044 |
| CKB | 100Kb | 6 (11.8%) | 2.00 (0.75- Inf) | 0.077 |
| CKB | 200Kb | 7 (13.7%) | 1.75 (0.78-7.00) | 0.091 |
| CKB | 500Kb | 10 (19.6%) | 1.43 (0.77-5.00) | 0.131 |
| UKB | 50Kb | 5 (9.8%) | 1.67 (0.71- Inf) | 0.093 |
| UKB | 100Kb | 5 (9.8%) | 1.67 (0.62- Inf) | 0.143 |
| UKB | 200Kb | 5 (9.8%) | 1.25 (0.56-5.00) | 0.277 |
| UKB | 500Kb | 9 (17.6%) | 1.29 (0.64-4.50) | 0.206 |
